# Progressive loss of conserved spike protein neutralizing antibody sites in Omicron sublineages is balanced by preserved T-cell recognition epitopes

**DOI:** 10.1101/2022.12.15.520569

**Authors:** Alexander Muik, Bonny Gaby Lui, Huitian Diao, Yunguan Fu, Maren Bacher, Aras Toker, Jessica Grosser, Orkun Ozhelvaci, Katharina Grikscheit, Sebastian Hoehl, Niko Kohmer, Yaniv Lustig, Gili Regev-Yochay, Sandra Ciesek, Karim Beguir, Asaf Poran, Özlem Türeci, Ugur Sahin

**Author notes:** Contributed equally (Alexander Muik, Bonny Gaby Lui).

## Abstract

The continued evolution of the SARS-CoV-2 Omicron variant has led to the emergence of numerous sublineages with different patterns of evasion from neutralizing antibodies. We investigated neutralizing activity in immune sera from individuals vaccinated with SARS-CoV-2 wild-type spike (S) glycoprotein-based COVID-19 mRNA vaccines after subsequent breakthrough infection with Omicron BA.1, BA.2, or BA.4/BA.5 to study antibody responses against sublineages of high relevance. We report that exposure of vaccinated individuals to infections with Omicron sublineages, and especially with BA.4/BA.5, results in a boost of Omicron BA.4.6, BF.7, BQ.1.1, and BA.2.75 neutralization, but does not efficiently boost neutralization of sublineages BA.2.75.2 and XBB. Accordingly, we found in *in silico* analyses that with occurrence of the Omicron lineage a large portion of neutralizing B-cell epitopes were lost, and that in Omicron BA.2.75.2 and XBB less than 12% of the wild-type strain epitopes are conserved. In contrast, HLA class I and class II presented T-cell epitopes in the S glycoprotein were highly conserved across the entire evolution of SARS-CoV-2 including Alpha, Beta, and Delta and Omicron sublineages, suggesting that CD8^+^ and CD4^+^ T-cell recognition of Omicron BQ.1.1, BA.2.75.2, and XBB may be largely intact. Our study suggests that while some Omicron sublineages effectively evade B-cell immunity by altering neutralizing antibody epitopes, S protein-specific T-cell immunity, due to the very nature of the polymorphic cell-mediated immune, response is likely to remain unimpacted and may continue to contribute to prevention or limitation of severe COVID-19 manifestation.

## Main Text

The SARS-CoV-2 Omicron variant of concern (VOC), that emerged in November 2021, contains over 30 amino acid alterations in its spike (S) glycoprotein as compared to the original Wuhan-Hu-1 (wild-type) strain that mediate partial escape from previously established immunity (*1–3*). Omicron sublineages BA.1, BA.2, BA.4, and BA.5 consecutively dominated the pandemic landscape. Immune escape became pronounced in the more recent sublineages, leading to the authorization of BA.1 and BA.4/BA.5 adapted vaccines (*4, 5*). While BA.5 has been the globally dominant sublineage since mid-2022, Omicron BA.2.75, BA.2.75.2, BA.4.6, BF.7, BQ.1.1 have locally increased, and XBB has displaced BA.5 in parts of Asia, whereas BQ.1.1 is currently becoming dominant over BA.5 in the United States and parts of Europe (*6–8*). Omicron BA.2.75 differs in five amino acids within the N-terminal domain from its parental sublineage BA.2 and previous Omicron VOCs (fig. S1). The receptor-binding domain (RBD) of BA.2.75 has three alterations that are not found in BA.2, of which G446S is shared with BA.1. BA.2.75.2 differs from BA.2.75 in the RBD alterations R346T and F486S. Omicron BA.4.6 and BF.7 are identical in their S glycoprotein sequence (called Omicron BA.4.6/BF.7 S in this study), which bears great similarity to the S glycoprotein sequence of their respective parental sublineages BA.4 and BA.5 (called Omicron BA.4/BA.5 S in this study due to sequence identity). A single R346T change within the RBD that distinguishes BA.4.6/BF.7 S glycoprotein from BA.4/BA.5 S glycoprotein abrogates neutralization by the therapeutic monoclonal antibody (mAb) cilgavimab (*9*). Cilgavimab in combination with tixagevimab (Evusheld™) is used for pre-exposure COVID-19 prophylaxis in immunocompromised patients. As tixagevimab lacks neutralizing activity against Omicron BA.4/BA.5 and their descendants, this antibody combination therapy is rendered fully ineffective against Omicron BA.4.6, BF.7 (*9, 10*) and BQ.1.1 (*11*), which additionally has a K444T alteration. The S protein of XBB, a recombinant strain (*12*), is altered at seven amino acid positions (four in NTD, three in RBD) relative to the main Omicron sublineages BA.1, BA.2, and BA.4/5 and also bears the R346T alteration. It is to be expected that virus evolution will continue. Real-time understanding of the transmissibility, pathogenicity, and immune evasion properties of new variants in conjunction with immunity patterns in the human population that are being shaped through repeated infections with different VOCs, vaccination, and booster cycles with different vaccines, will continue to be critical to assessing the level of risk to public health going forward.

In the face of this highly dynamic situation, we have set up a COVID-19 pandemic preparedness and rapid response strategy. Our approach includes the screening of emerging and circulating variants through an artificial intelligence/machine learning-based early warning system (*13*), testing prototypical mRNA vaccines adapted to these variants in mouse studies (*16*), and mapping SARS-CoV-2 T-cell epitopes recognized by the human T-cell repertoire (*19, 20*). Further, studying the neutralizing antibody activity of individuals following breakthrough infections with the latest circulating variants allows us to detect potential immune escape patterns early and informs on the need for rapid vaccine adaptation strategies (*14–18*). In continuation of this work, the current study assessed Omicron sublineages BA.4.6, BF.7, BA.2.75, BA.2.75.2, BQ.1.1, and XBB, as these sublineages are currently establishing dominance over BA.5.

We investigated neutralizing activity of immune sera from individuals who received three or four doses of SARS-CoV-2 wild-type S glycoprotein-based mRNA COVID-19 vaccines (BNT162b2/mRNA-1273 homologous or heterologous regimens) with or without subsequent breakthrough infection by different Omicron sublineages. Specifically, the following cohorts were investigated: BNT162b2 triple-vaccinated SARS-CoV-2-naïve individuals (BNT162b2^3^; n=18; age <55 years), BNT162b2 quadruple-vaccinated SARS-CoV-2-naïve elderly (>60 years) individuals (BNT162b2^4^; n=15), and triple mRNA vaccinated individuals who experienced breakthrough infection with Omicron BA.1 (mRNA-Vax^3^ + BA.1; n=14), BA.2 (mRNA-Vax^3^ + BA.2, n=19), or BA.4/BA.5 (mRNA-Vax^3^ + BA.4/BA.5, n=17) (fig. S2). As a fourth vaccine dose is recommended for the elderly and given the high frequency of breakthrough infections with Omicron sublineages we considered this study population to be representative for a large proportion of the European and North American population. Serum neutralizing activity was tested in a well-characterized pseudovirus neutralization test (pVNT) (*14–16*) by determining 50% pseudovirus neutralization (pVN_50_) geometric mean titers (GMTs) for pseudoviruses bearing the S glycoproteins of the SARS-CoV-2 wild-type strain or of Omicron sublineages.

In the triple-/quadruple-vaccinated individuals without breakthrough infection, pVN_50_ GMTs against Omicron BA.4/BA.5 were 5 to 6-fold lower than GMTs against the wild-type strain (GMTs against BA.4/BA.5 were 69 and121, respectively) (Fig. 1a). GMTs against BA.4/BA.5 were similarly reduced in BA.1 convalescents (GMT 263, 5-fold lower than wild-type), whereas in the BA.2 and BA.4/BA.5 convalescent cohorts, titers against BA.4/BA.5 remained higher (GMTs of 386 and 521, respectively; 3- and 2-fold lower than wild-type).

**Fig. 1.**
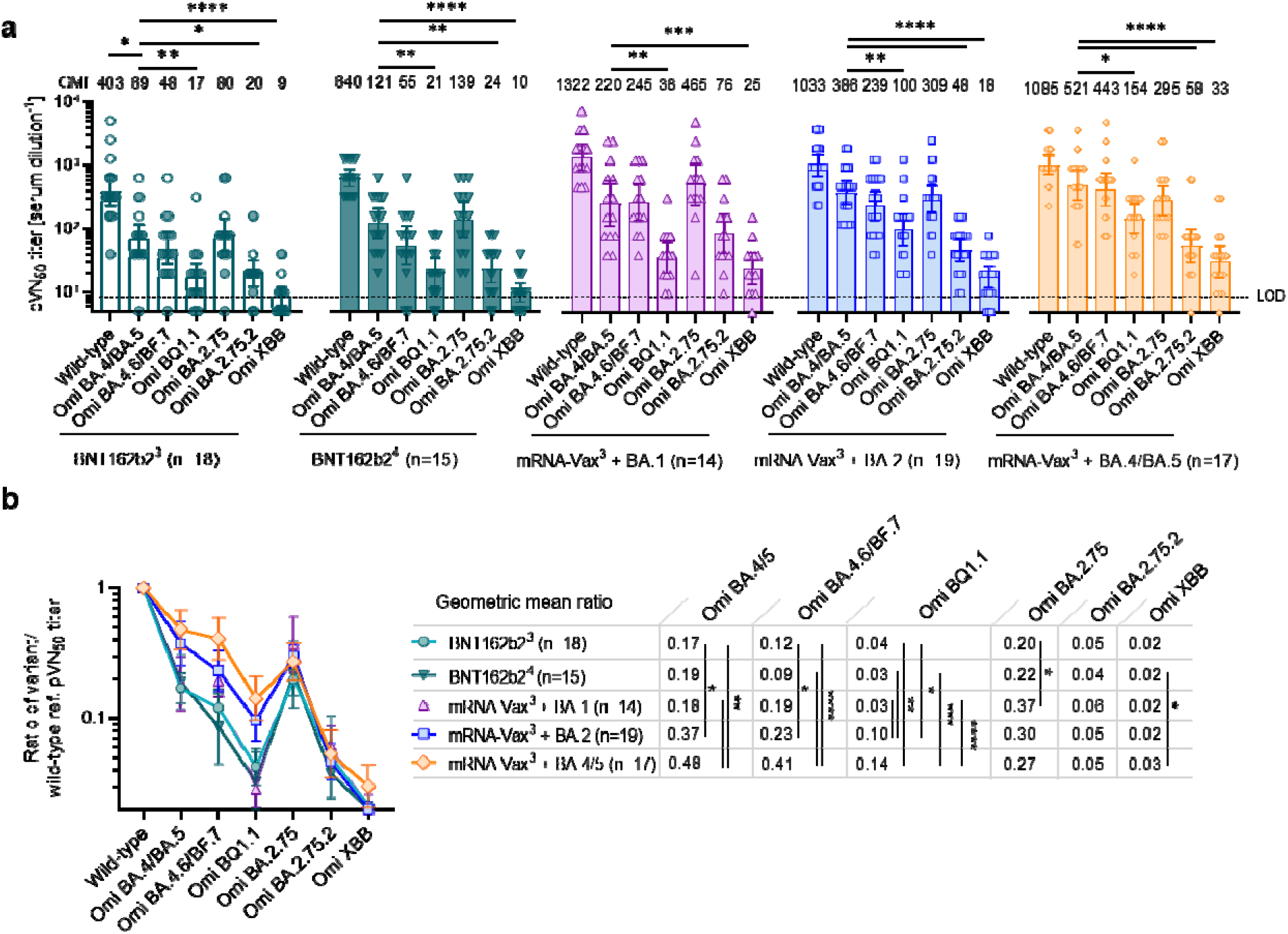
Distinct cross-neutralization of Omicron sublineages by vaccine-elicited and convalescent human immune sera.

In all three convalescent cohorts, neutralizing titers against Omicron BA.4.6/BF.7 and BA.2.75 were robustly above those of triple-/quadruple-vaccinated SARS-CoV-2 naïve individuals (GMT range 239-525 for convalescents as compared to 55-139 for naïves). In the convalescents, Omicron BA.4.6/BF.7 and BA.2.75 GMTs were largely comparable with no significant differences to those against Omicron BA.4/BA.5. In contrast, pVN_50_ titers against Omicron BQ.1.1, BA.2.75.2, and XBB were significantly lower than those against BA.4/BA.5 across cohorts. Titers against BQ.1.1 were overall very low in the SARS-CoV-2 naïve vaccinated cohorts and BA.1 convalescents (GMTs ≤38) and moderately higher in the BA.2 and BA.4/BA.5 convalescent cohorts (GMTs 100 and 154, respectively). Titers against BA.2.75.2 and XBB were low across cohorts (GMTs ≤88 and ≤33, respectively).

To assess neutralization breadth irrespective of the magnitude of antibody titers we normalized the Omicron sublineage pVN_50_ GMTs against those for the wild-type strain. GMT ratios for all Omicron subvariant pseudoviruses were comparable between the BNT162b2^3^ and BNT162b2^4^ cohorts (Fig. 1b). Hence, both neutralizing antibody titers and variant cross-neutralization were broadly similar in quadruple-vaccinated elderly individuals and triple-vaccinated younger individuals. GMT ratios were in the range of 0.09-0.22 for BA.4/BA.5, BA.4.6/BF.7, and BA.2.75 and ≤0.05 for BQ.1.1, BA.2.75.2, and XBB in both cohorts.

Cross-neutralization of BA.4/BA.5 and BA.4.6/BF.7 was significantly (p<0.05) higher in sera from BA.2.convalescents as compared to triple-vaccinated individuals (GMT ratios 0.37 vs 0.17 for BA.4/BA.5, and 0.23 vs 0.12 for BA.4.6/BF.7) and even more so in BA.4/BA.5-convalescents for both the BA.4/BA.5 pseudovirus (GMT ratio 0.48, p<0.01 versus BNT162b2^3^) and the BA.4.6/BF.7 pseudovirus (GMT ratio 0.41, p<0.0001). Cross-neutralization of BA.4.6/BF.7 was also significantly (p<0.0001) stronger in BA.4/BA.5 convalescents compared to quadruple-vaccinated individuals. While BQ.1.1 was cross-neutralized less efficiently than BA.4/BA.5 in all cohorts (GMT ratios ≤0.14), cross-neutralization in BA.4/BA.5 and BA.2 convalescents remained significantly stronger compared to SARS-CoV-2 naïve triple or quadruple-vaccinated and BA.1 breakthrough infected cohorts. Cross-neutralization of BA.2.75, BA.2.75.2, and XBB pseudoviruses was broadly comparable across cohorts. GMT ratios were in the range of 0.20-0.37 for BA.2.75 and ≤0.06 for BA.2.75.2 and XBB. Together these data show that partial neutralization of some Omicron sublineages is retained, being broadest in BA.4/BA.5 convalescent individuals. In contrast, sublineages BA.2.75.2 and XBB have evolved to largely evade neutralizing antibody responses in vaccinated individuals and in those with breakthrough infections with previous and currently circulating Omicron sublineages.

We next sought to determine whether distinct cross-neutralization of individual SARS-CoV-2 variants is reflected by the degree of B-cell epitope conservation with respect to the wild-type strain. For this, we analyzed a total of 506 neutralizing B-cell epitopes within the S glycoprotein NTD and RBD, consisting of experimentally confirmed epitopes from the Immune Epitope Database (IEDB) and epitopes computationally deduced from structures of SARS-CoV-2 neutralizing antibodies from the Coronavirus Antibody Database (CoV-AbDab). Of these 506 epitopes, 462 (91%) included a position that was altered in at least one of the analyzed variants.

We found that B-cell epitopes were partially conserved in the earlier variants Alpha, Beta, and Delta (≥43%) (Fig 2, Fig S3, and Table S2), whereas with the occurrence of the heavily RBD-mutated Omicron BA. 1 variant most of these epitopes were altered and remained so for the Omicron lineage (≤22% conservation). This was particularly the case in BA.2.75.2 and XBB (≤12% conservation), in line with our cross-neutralization data.

**Fig. 2.**
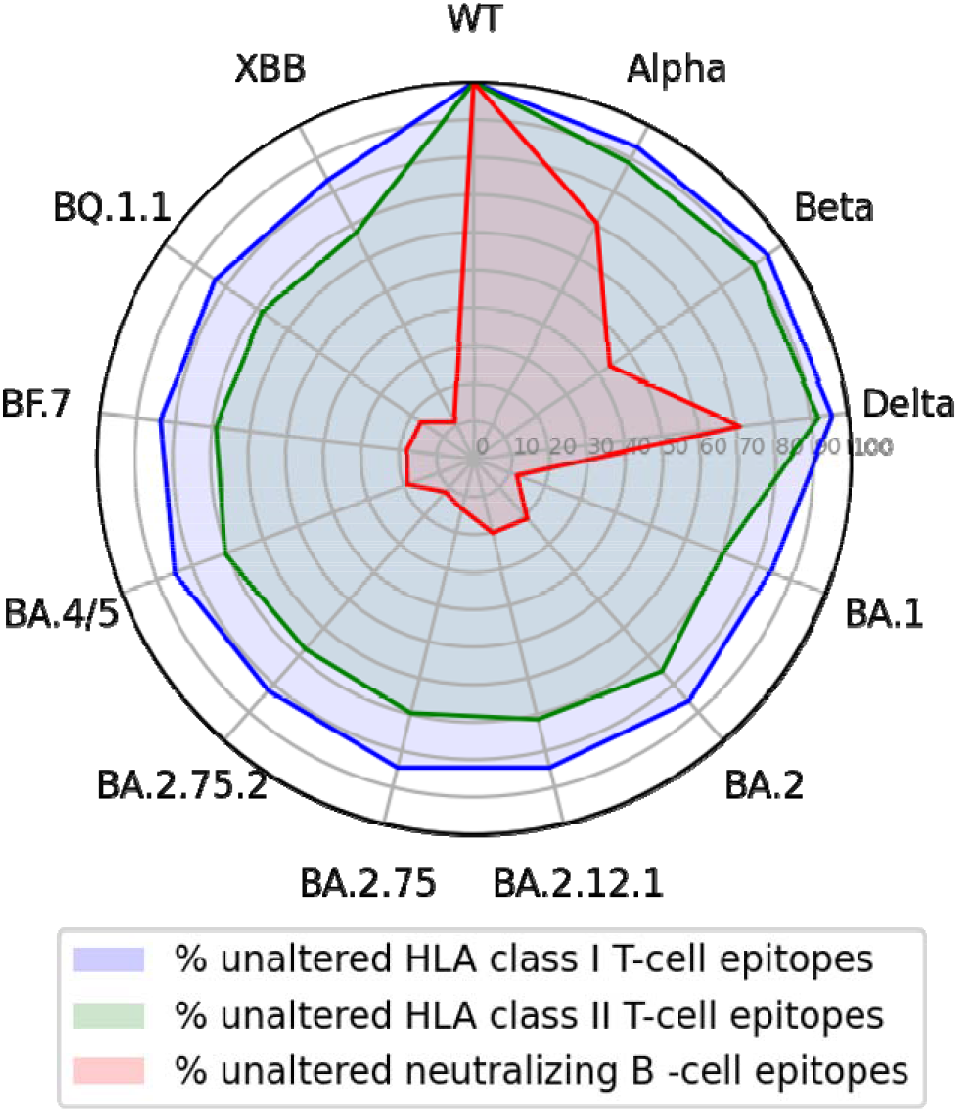
T-cell epitopes of the wild-type SARS-CoV-2 S glycoprotein but not epitopes for neutralizing antibodies are largely conserved across variants.

Neutralizing antibodies are an important component of immunity against SARS-CoV-2 yet not the only one (*17*). T cells are known to contribute to the adaptive anti-viral immune response. Unlike antibodies, which do not require HLA presentation and often depend on specific epitope conformations, T-cell recognition of the viral proteins is distributed across varying linear epitopes in different individuals based on their specific composition of HLA class I and II alleles. This renders T-cell immunity in the population harder to evade through accumulation of mutations. Cytotoxic CD8^+^ T cells that recognize HLA class I presented epitopes play a critical role in susceptibility to severe COVID-19 disease, hospitalization and death (*18, 19*). HLA class II dependent CD4^+^ helper T cells typically orchestrate the immune response by their pleiotropic functions and are also considered to have an important role in the prevention of severe disease (*20, 21*).

To estimate the impact of SARS-CoV-2 evolution on T-cell immunity, we assessed the degree of conservation of T-cell epitopes localized in the S glycoproteins of various VOCs. To this aim we filtered experimentally confirmed SARS-CoV-2 epitopes reported for HLA class I and II alleles from the IEDB database. In total, 594 and 559 unique epitopes were collected for HLA class I and class II, respectively. For stringency, in this analysis we only retained experimentally confirmed minimal epitopes of either HLA class I or HLA class II in a human host and removed any inferred or predicted entries. Epitopes were also filtered by length (8-14 for HLA class I and 12-20 for HLA class II). Following filtering, 260 unique HLA class I epitope sequences remained, of which 244 were found in the wild-type strain S glycoprotein. Of these, 71 epitopes (27.3%) included a position reported to be mutated in at least one analyzed variant. Of 468 HLA class II epitopes, 230 (49%) were observed to cover region that contains mutation in at least one analyzed variant. HLA class I and class II epitopes post filtering covered in total 40 unique alleles for both class I and class II.

Approximately 90% of CD8^+^ and CD4^+^ T-cell epitopes of the wild-type S glycoprotein were fully conserved in the Alpha, Beta, and Delta variants (Fig 2, fig S3, Tables S3 and S4) and over 80% of CD8^+^ and ~70% CD4^+^ T-cell epitopes were fully conserved in Omicron sublineages including BA.2.75.2, BQ.1.1, and XBB (Fig. 2 and fig. S3), suggesting that T-cell responses against Omicron sublineages may remain largely intact in individuals immunized with wild-type strain-based vaccines.

In summary, our findings provide insights into two mechanisms: firstly, the neutralizing activity of human sera in the context of currently circulating VOCs, and second, the potential of the SARS-CoV-2 S glycoprotein directed CD8^+^ and CD4^+^ T-cell repertoire.

With regard to humoral immunity, we show that exposure of vaccinated individuals to Omicron BA.4/BA.5 refocuses neutralizing antibody responses towards neutralization of BA.4/BA.5 itself, that goes along with partial cross-neutralization of BA.4.6/BF.7, BA.2.75 and at a lower degree of BQ.1.1 but displays poor activity against Omicron BA.2.75.2 and XBB.

That the Omicron BA.4.6/BF.7-neutralizing activity and even more so neutralization of BQ.1.1 by sera from vaccinated individuals and BA.2 convalescents is further reduced as compared to their activity against BA.4/BA.5 suggests that the R346T and the N460K alterations mediate further escape from neutralizing antibodies in human sera. Convergent evolution of the RBD at these critical residues in Omicron BA.4.6, BF.7, BQ.1.1, BA.2.75.2 and XBB (*7, 12*) suggests that the resulting immune evasion may confer a growth advantage. Reduced neutralization of BA.2.75.2 as compared to BA.2.75 confirms a prominent role of R346T as well as of alterations at F486 in escape from neutralizing antibodies (*22*).

Consistent with previous reports we show that cross-neutralization of Omicron BA.2.75 and BA.4/BA.5 are broadly comparable (*10, 23, 24*) indicating factors other than immune evasion to be involved in growth advantage of BA.2.75 over BA.5. Minor differences in the sensitivity of Omicron BA.2.75 and BA.4/BA.5 to neutralization by BA.1- and BA.4/BA.5-convalescent sera may indicate amino acid changes with a context-dependent role in immune evasion.

Our findings are supportive for suitability of Omicron BA.4/BA.5-adapted vaccines to boost neutralizing activity against several circulating or emerging variants of potential relevance, such as BA.4.6 and BF.7. They also show that Omicron sublineages continue to accumulate mutations that disrupt critical B-cell neutralization epitopes, and that further boosters or variant adaptations may be required in future.

With regard to cell-mediated immunity, we show that HLA class I and class II presented T-cell epitopes remained mostly unaltered across the evolution of SARS-CoV-2 including Omicron sublineages, suggesting that CD8^+^ and CD4^+^ T-cell immunity against Omicron BQ.1.1, BA.2.75.2, and XBB may be largely intact, despite profound neutralizing antibody evasion. This fits in with previous reports, e.g. describing the existence of degenerate T-cell epitopes located within conserved S protein regions (*25*) and showing that wildtype-strain-vaccinated individuals retain T-cell immunity against Omicron BA.1 (*26–28*). Indeed, a fundamental difference of T-cell versus B-cell mediated immunity is that owing to the highly polymorphic nature of HLA molecules (*29–31*), the T-cell mediated layer of immunity is more robust against population-level breaches by VOCs. Our observations indicate that T-cell immunity may mitigate the lack of neutralizing antibody activity in preventing or limiting severe COVID-19 and further encourages development of vaccine formats that induce functional and broad SARS-CoV-2-directed CD8^+^ T-cell immunity concurrently to boosting antibody responses.

Cohorts and serum sampling as described in fig. S2. (a) 50% pseudovirus neutralization (pVN_50_) geometric mean titers (GMTs) against the indicated SARS-CoV-2 wild-type strain or Omicron variants of concern (VOCs). Values above bar graphs represent group GMTs. For titer values below the limit of detection (LOD), LOD/2 values were plotted. The non-parametric Friedman test with Dunn’s multiple comparisons correction was used to compare neutralizing titers against the Omicron BA.4/BA.5 pseudovirus (which represents currently dominating BA.5) with titers against the other pseudoviruses. Multiplicity-adjusted p values are shown. (b) SARS-CoV-2 VOC pVN_50_ GMTs normalized against the wild-type strain pVN_50_ GMT (ratio VOC to wildtype). Group geometric mean ratios with 95% confidence intervals are shown. The non-parametric Kruskal-Wallis test with Dunn’s multiple comparisons correction was used to compare the VOC GMT ratios between cohorts. ****, P<0.0001; ***, p<0.001; **, P<0.01; *, P<0.05. Serum was tested in duplicate.

Percentages of unaltered neutralizing B-cell epitopes (restricted to NTD and RBD) present in each variant strain compared to SARS-CoV-2 wild-type are represented by the red shaded area. Percentages of unaltered S glycoprotein linear HLA class I and II T-cell epitopes present in each variant strain as compared to SARS-CoV-2 S wild-type are shown in blue (HLA class I) and green (HLA class II) shaded area. The B-cell epitopes were either retrieved from IEDB on November 25, 2022 or calculated from resolved antigen-antibody protein structures deposited in the Coronavirus antibody database (CoV-AbDab) (*32*) using a similar computational method to the one used in our early warning system (*13*). T-cell epitopes were retrieved from the Immune Epitope Database (IEDB) on November 11, 2022.

## Acknowledgments

We thank the BioNTech global clinical trial participants (NCT05004181, NCT04955626), and the Omicron BA.1, BA.2 and BA.4/5 convalescent research study participants from whom the post-immunization human sera were obtained. We thank the many colleagues at BioNTech and Pfizer who developed and produced the BNT162b2 vaccine candidate. We thank Sabrina Jägle and Nina Beckmann for logistical support. We thank Svetlana Shpyro, Sayeed Nadim, Christina Heiser, Ayca Telorman, Claudia Müller, Amy Wanamaker, Nicki Williams, and Jennifer VanCamp for sample demographics support.

## Author contributions

U.S., Ö.T., and A.M. conceived and conceptualized the work. A.M., A.P., K.B., and B.G.L. planned and supervised experiments and epitope analyses. G.R.Y., O.O., K.G., J.G., N.K., S.H. and S.C. coordinated and conducted sample collection. K.G., Y.L., and G.R.Y. coordinated sample shipments and clinical data transfer. B.G.L., and M.B. performed experiments. A.M., B.G.L., A.P, Y.F., and H.D. analyzed data. U.S., Ö.T., A.M., A.P., A.T., and B.G.L. interpreted data and wrote the manuscript. All authors supported the review of the manuscript.

## Data and materials availability

Materials are available from the authors under a material transfer agreement with BioNTech.

## Supplementary materials

Materials and Methods

Figs. S1-S4

Tables S1-S9

