## Supplement for "Progressive loss of conserved spike protein neutralizing antibody sites in Omicron sublineages is balanced by preserved T-cell recognition epitopes"

**This PDF file includes:**

Materials and Methods

Fig. S1 to S4

Tables S1 and S5 to S9

**Supplied as separate files:**

Tables S2 to S4

**Materials and Methods**

##### **Study design, recruitment of participants and sample collection**

The objective was to investigate the capability of five different human sera panels to cross-neutralize Omicron BA.4.6/BF.7 and BA.2.75 sub-lineages in comparison to SARS-CoV-2 wild-type and Omicron BA.4/BA.5. The sera panels were derived from (i) SARS-CoV-2-naïve, triple-BNT162b2-vaccinated adults < 55years of age (BNT162b2^3^), (ii) SARS-CoV-2-naïve quadruple-BNT162b2-vaccinated adults >60 years of age (BNT162b2^4^), and triple-mRNA vaccinated individuals (BNT162b2 or mRNA-1273) with a confirmed subsequent SARS-CoV-2 breakthrough infection which either occurred in a period of dominance of (iii) the Omicron BA.1 lineage (November 2021 to mid-January 2022; mRNA-Vax^3^ + BA.1) (*1*), (iv) Omicron BA.2 lineage (March to May 2022; mRNA-Vax^3^ + BA.2) (*2*) or (v) Omicron BA.4/BA.5 lineage in Germany (mid-June to mid-July 2022; mRNA-Vax^3^ + BA.4/5) (*3*). Serum neutralizing capability was characterized using pseudovirus neutralization assays. SARS-CoV-2 wild-type and Omicron BA.4/BA.5 neutralization data for cohorts BNT162b2^3^, mRNA-Vax^3^ + BA.1, mRNA-Vax^3^ + BA.2, mRNA-Vax^3^ + BA.4/5 were previously published (*1-3*).

Participants from the mRNA-Vax^3^ + Omi BA.1, mRNA-Vax^3^ + Omi BA.2, and mRNA-Vax^3^ + BA.4/5 cohorts were recruited from University Hospital, Goethe University Frankfurt as part of a non-interventional study (protocol approved by the Ethics Board of the University Hospital [No. 2021-560]) researching patients that had experienced Omicron breakthrough infection following vaccination for COVID-19. Individuals from the BNT162b2^3^ and BNT162b2^4^ cohort were consented participants in the Phase 2 trial BNT162-17 (NCT05004181) and BNT162-16 Substudy F (NCT04955626), respectively. All participants had no documented history of SARS-CoV-2 infection prior to vaccination. Participants were free of symptoms at the time of blood collection.

Serum was isolated by centrifugation of drawn blood at 2000 x g for 10 minutes and cryopreserved until use.

##### **VSV-SARS-CoV-2 S variant pseudovirus generation**

A recombinant replication-deficient vesicular stomatitis virus (VSV) vector that encodes green fluorescent protein (GFP) and luciferase instead of the VSV-glycoprotein (VSV-G) was pseudotyped with SARS-CoV-1 S glycoprotein (UniProt Ref: P59594) or with SARS-CoV-2 S glycoprotein derived from either the wild-type strain (Wuhan-Hu-1, NCBI Ref: 43740568), the Omicron BA.4/BA.5 variant (alterations: T19I, Δ24-26, A27S, Δ69/70, G142D, V213G, G339D, S371F, S373P, S375F, T376A, D405N, R408S, K417N, N440K, L452R, S477N, T478K, E484A, F486V, Q498R, N501Y, Y505H, D614G, H655Y, N679K, P681H, N764K, D796Y, Q954H, N969K), the Omicron BA.4.6/BF.7 variant (alterations: T19I, Δ24-26, A27S, Δ69/70, G142D, V213G, G339D, R346T, S371F, S373P, S375F, T376A, D405N, R408S, K417N, N440K, L452R, S477N, T478K, E484A, F486V, Q498R, N501Y, Y505H, D614G, H655Y, N679K, P681H, N764K, D796Y, Q954H, N969K), the Omicron BQ.1.1 variant (alterations: T19I, Δ24-26, A27S, Δ69/70, G142D, V213G, G339D, R346T, S371F, S373P, S375F, T376A, D405N, R408S, K417N, N440K, K444T, L452R, S477N, T478K, E484A, F486V, Q498R, N501Y, Y505H, D614G, H655Y, N679K, P681H, N764K, D796Y, Q954H, N969K), the Omicron BA.2.75 variant (alterations: T19I, Δ24-26, A27S, G142D, K147E, W152R, F157L, I210V, V213G, G257S, G339H, S371F, S373P, S375F, T376A, D405N, R408S, K417N, N440K, G446S, N460K, S477N, T478K, E484A, Q498R, N501Y, Y505H, D614G, H655Y, N679K, P681H, N764K, D796Y, Q954H, N969K), the Omicron BA.2.75.2 variant (alterations: T19I, Δ24-26, A27S, G142D, K147E, W152R, F157L, I210V, V213G, G257S, G339H, R346T, S371F, S373P, S375F, T376A, D405N, R408S, K417N, N440K, G446S, N460K, S477N, T478K, E484A, F486S, Q498R, N501Y, Y505H, D614G, H655Y, N679K, P681H, N764K, D796Y, Q954H, N969K, D1199N), and the Omicron XBB variant (alterations: T19I, Δ24-26, A27S, V83A, G142D, Δ144, H146Q, Q183E, V213E, G339H, R346T, L368I, S371F, S373P, S375F, T376A, D405N, R408S, K417N, N440K, V445P, G446S, N460K, S477N, T478K, E484A, F486S, F490S, Q498R, N501Y, Y505H, D614G, H655Y, N679K, P681H, N764K, D796Y, Q954H, N969K) according to published pseudotyping protocols (*4*). A diagram of SARS-CoV-2 S glycoprotein alterations is shown in fig. S4 and a separate alignment of S glycoprotein alterations in Omicron VOCs is displayed in fig. S1.

In brief, HEK293T/17 monolayers (ATCC® CRL-11268™) cultured in Dulbecco’s modified Eagle’s medium (DMEM) with GlutaMAX™ (Gibco) supplemented with 10% heat-inactivated fetal bovine serum (FBS [Sigma-Aldrich]) (referred to as medium) were transfected with Sanger sequencing-verified variant-specific SARS-CoV-2 S expression plasmid with Lipofectamine LTX (Life Technologies) following the manufacturer’s instructions. At 24 hours after transfection, the cells were infected at a multiplicity of infection (MOI) of three with VSV-G complemented VSVΔG vector. After incubation for 2 hours at 37 °C with 7.5% CO_2_, cells were washed twice with phosphate buffered saline (PBS) before medium supplemented with anti-VSV-G antibody (clone 8G5F11, Kerafast Inc.) was added to neutralize residual VSV-G-complemented input virus. VSV-SARS-CoV-2-S pseudotype-containing medium was harvested 20 hours after inoculation, passed through a 0.2 µm filter (Nalgene) and stored at -80 °C. The pseudovirus batches were titrated on Vero 76 cells (ATCC® CRL-1587™) cultured in medium. The relative luciferase units induced by a defined volume of a SARS-CoV-2 wild-type strain S glycoprotein pseudovirus reference batch previously described in Muik et al., 2021 (*5*), that corresponds to an infectious titer of 200 transducing units (TU) per mL, was used as a comparator. Input volumes for the SARS-CoV-2 variant pseudovirus batches were calculated to normalize the infectious titer based on the relative luciferase units relative to the reference.

##### **Pseudovirus neutralization assay**

We previously showed that VOC neutralizing titers detected in the pseudovirus neutralization assay strongly corelated with those from authentic live SARS-CoV-2 virus neutralization assays (*6*). Vero 76 cells were seeded in 96-well white, flat-bottom plates (Thermo Scientific) at 40,000 cells/well in medium 4 hours prior to the assay and cultured at 37 °C with 7.5% CO_2_. Human serum samples were 2-fold serially diluted in medium with dilutions ranging from 1:10 to 1:10,240. VSV-SARS-CoV-2-S particles were diluted in medium to obtain 200 TU in the assay. Serum dilutions were mixed 1:1 with pseudovirus (n=2 technical replicates per serum per pseudovirus) for 30 minutes at room temperature before being added to Vero 76 cell monolayers and incubated at 37°C with 7.5% CO_2_ for 24 hours. Supernatants were removed and the cells were lysed with luciferase reagent (Promega). Luminescence was recorded on a CLARIOstar® Plus microplate reader (BMG Labtech), and neutralization titers were calculated as the reciprocal of the highest serum dilution that still resulted in 50% reduction in luminescence. Results for all pseudovirus neutralization experiments were expressed as geometric mean titers (GMT) of duplicates. If no neutralization was observed, an arbitrary titer value of half of the limit of detection [LOD] was reported. Neutralization titers in human sera are shown in Tables S4 to S8.

##### **T-cell epitope conservation analysis**

To estimate the rate of non-synonymous mutations in T cell epitopes in the spike glycoprotein, we obtained CD4^+^ and CD8^+^ T cell epitopes confirmed via experimental assays which were reported in the Immune Epitope Database (<https://www.iedb.org/>) (*7*) . The experimental assays confirming the reactivity of these epitopes relied on multimer analysis, ELISpot or ELISpot-like assays, T cell activation assays, etc.

The database was filtered using the following criteria: Organism: SARS-COV2; Antigen: Spike glycoprotein; Positive Assay; No B cell assays; No MHC assays; MHC Restriction Type: Class I (or II); Host: Homo sapiens (human). In order to restrict the dataset to confirmed minimal epitopes only, the resulting tables were filtered by removing entries that were deduced from a reactive overlapping peptide pool or relied on prediction to identify the minimal epitope. Only epitopes of length 8-14 amino acids (for HLA-I) or 12-20 amino acids (for HLA-II) were retained.

##### **B-cell epitope conservation analysis**

The B cell epitopes are collected from Immune Epitope Database (IEDB, <https://www.iedb.org/>) and Coronavirus Antibody Database (CoV-AbDab), <http://opig.stats.ox.ac.uk/webapps/covabdab/>) (*8*). The IEDB database was queried on November 25, 2022 using the following criteria: Organism: SARS-COV2; Antigen: Spike glycoprotein; Positive Assay; No T cell assays; No MHC assays; Host: Homo sapiens (human); B Cell Assays: neutralization | biological activity (neutralization). The resulting table was filtered by limiting epitopes to NTD or RBD targeting with a minimum number of 3 amino acids. Of the 430 unique epitope sequences obtained in this approach from IEDB, 355 were found in the wild-type strain S glycoprotein. Protein data bank (PDB) structures and neutralization information were collected from CoV-AbDab (updated on October 3, 2022). The epitopes were calculated from the structures corresponding to SARS-CoV-2 neutralizing antibodies with human hosts, in a similar method to the one used in the Early Warning System (*9*). Precisely, given one or multiple antigen chains, the epitope consists of positions on the antigen chains that are in contact with antibody heavy or light chains. Two amino acids are in contact if the smallest Euclidean distance between their atoms was smaller than 4 Ångstroms and the largest Euclidean distance was smaller than 20 Ångstroms. The resulting epitopes were further limited to NTD or RBD targeting with a minimum number of 3 amino acids. Of the 295 unique epitope sequences obtained in this approach from CoV-AbDab, 234 were found in the wild-type strain spike glycoprotein. Merging structure-based epitopes and IEDB retrieved epitopes resulted in 506 unique epitopes. Of these, 462 epitopes (91%) included a position reported to be mutated in at least one of the variants investigated in our study.

##### **Statistical analysis**

The statistical method of aggregation used for the analysis of antibody titers is the geometric mean and for the ratio of SARS-CoV-2 VOC titer and wild-type strain titer the geometric mean and the corresponding 95% confidence interval. The use of the geometric mean accounts for the non-normal distribution of antibody titers, which span several orders of magnitude. The Friedman test with Dunn’s correction for multiple comparisons was used to conduct pairwise signed-rank tests of group geometric mean neutralizing antibody titers with a common control group. The Kruskal-Wallis test with Dunn’s correction for multiple comparisons was used to conduct unpaired signed-rank tests of group GMT ratios. All statistical analyses were performed using GraphPad Prism software version 9.

**fig. S1**

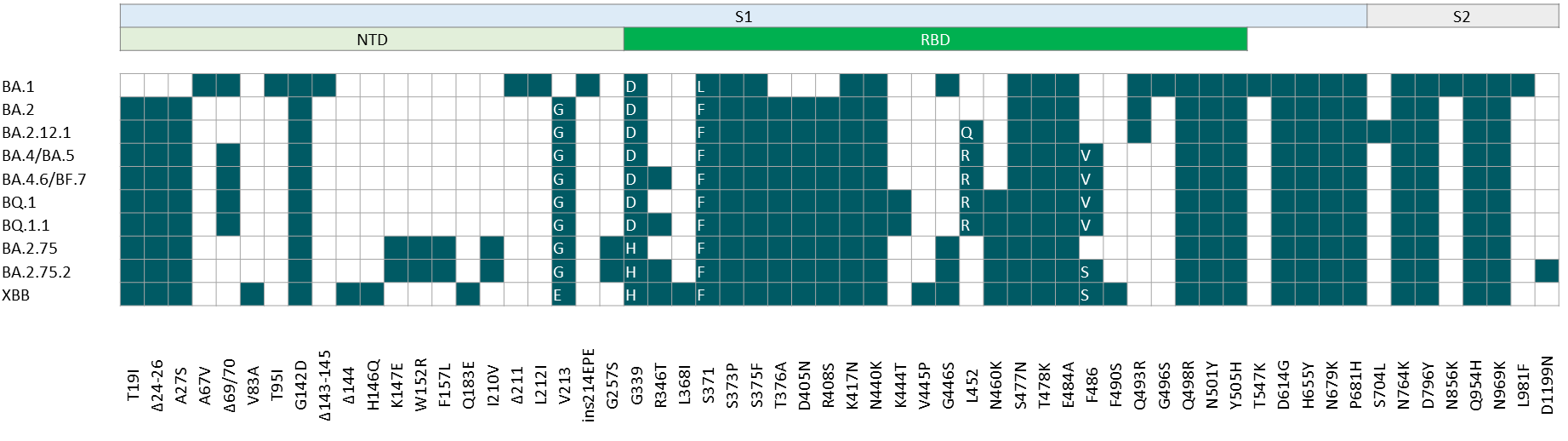

**Fig. S1. Alterations of the spike glycoprotein amino acid sequence of SARS-CoV-2 Omicron sub-lineages.**

White letters in boxes indicate the amino acid substitution per sub-lineage; Δ, deletion; ins, insertion; S1, S1-subunit of the S glycoprotein; S2, S2-subunit of the S glycoprotein; NTD, N-terminal domain; RBD, receptor-binding domain

**fig. S2**

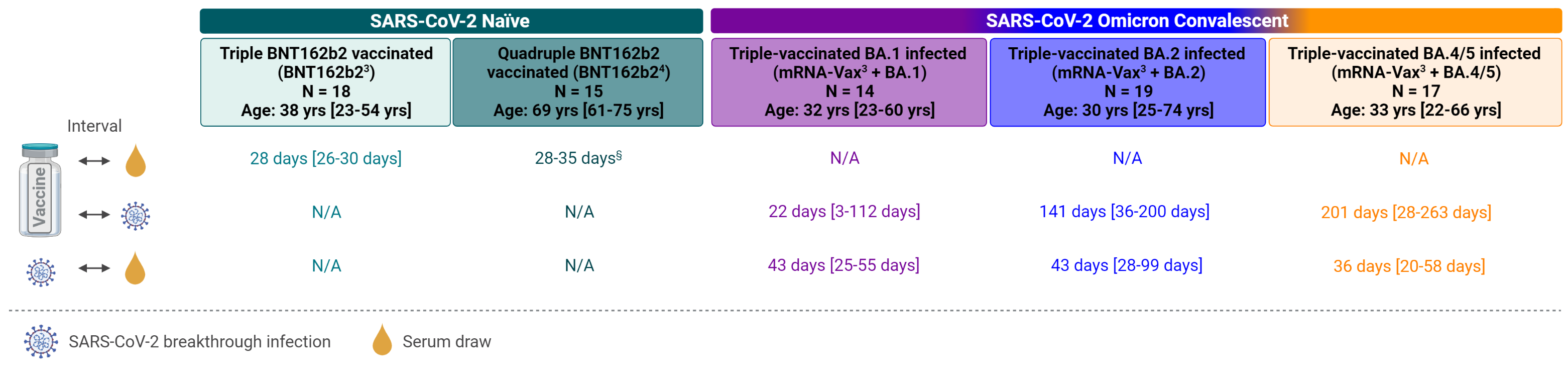

**Fig. S2. Cohorts and sampling**

Serum samples were drawn from five cohorts: SARS-CoV-2-naïve individuals triple-vaccinated with BNT162b2 (BNT162b2^3^, light-green) or quadruple-vaccinated with BNT162b2 (BNT162b2^4^, dark-green), and individuals with three doses of mRNA COVID-19 vaccine (BNT162b2/mRNA-1273 homologous or heterologous regimens) who subsequently had a breakthrough infection with Omicron BA.1 (mRNA-Vax3 + BA.1, purple), with BA.2 (mRNA-Vax3 + BA.2, blue) or with BA.4/BA.5 (mRNA-Vax3 + BA.4/5, orange). Breakthrough infections occurred at a time of respective VOC dominance (BA.1: November 2021 to January 2021, BA.2: March to May 2022, BA.4/5: mid-June to mid-July 2022) and/or were variant confirmed by genome sequencing. For convalescent cohorts, relevant intervals between key events such as the most recent vaccination, SARS-CoV-2 infection, and serum isolation are indicated. All values specified as median-range. The age/gender composition of the cohorts is further detailed in Table S1. Data for the cohorts BNT162b2^3^, mRNA-Vax^3^ + BA.1, mRNA-Vax^3^ + BA.2, and mRNA-Vax^3^ + BA.4/5 were previously published (*1, 2*){Muik, 2022 #88.

N/A, not applicable; ^§^, Serum draw was performed between 28 to 35 Days after vaccination as per protocol; Schematic was created with BioRender.com

**fig. S3**

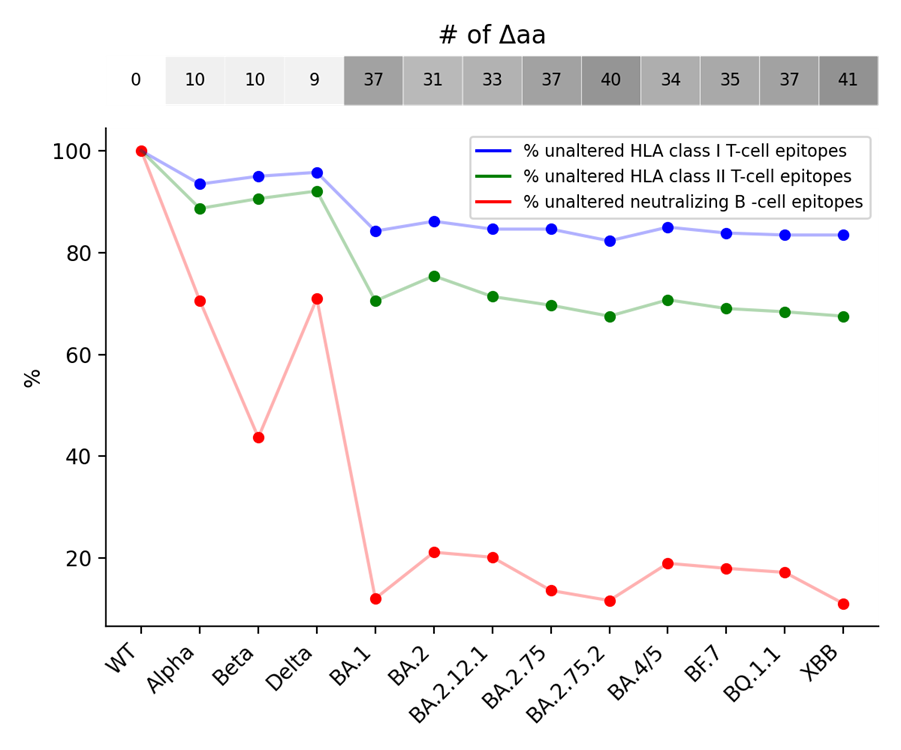

**Fig. S3. T-cell epitopes of the wild-type SARS-CoV-2 S glycoprotein but not epitopes for neutralizing antibodies are largely conserved in Omicron sublineages**

Percentages of unaltered neutralizing B-cell epitopes localized in NTD and RBD present in each variant strain compared to SARS-CoV-2 wild-type (red) and of unaltered S glycoprotein linear T-cell epitopes (Blue for HLA class I, green for HLA class II) are shown. The B-cell epitopes were either retrieved from IEDB on November 25, 2022 or calculated from resolved antigen-antibody protein structures deposited in the Coronavirus antibody database (CoV-AbDab) (*8*) using a similar computational method to the one used in our Early Warning System (*9*). T cell epitopes were retrieved from the Immune Epitope Database (IEDB) on November 11, 2022. Numbers above the graph depict numbers of amino acid alterations found in each variant relative to the wild-type strain.

**Fig. S4**

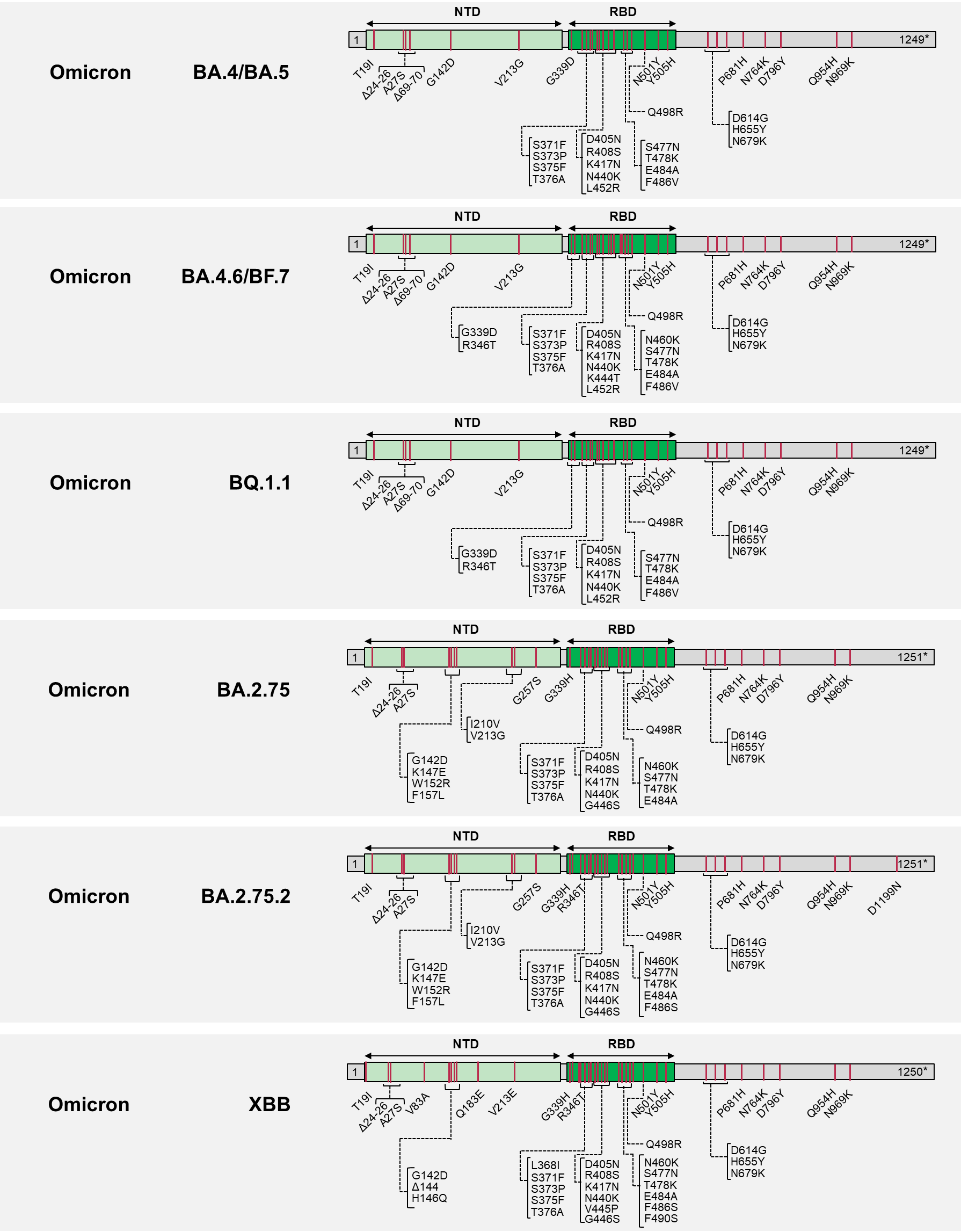

### **Fig. S4. Characterization of SARS-CoV-2 S glycoproteins used in the VSV-SARS-CoV-2 variant pseudovirus neutralization assays.**

### The sequence of the Wuhan-Hu-1 isolate SARS-CoV-2 S glycoprotein (GenBank: QHD43416.1) was used as reference. Amino acid positions, amino acid descriptions (one letter code) and kind of alterations (substitutions, deletions, insertions) are indicated. NTD, N-terminal domain; RBD, Receptor-binding domain, Δ, deletion; ins, insertion; *, Cytoplasmic domain truncated for the C-terminal 19 amino acids.

**Table S1.** **Vaccinated individuals analyzed for neutralizing antibody responses.**

| **Characteristic** | **BNT162b2^3^**  **(n=18)** | **BNT162b2^4^**  **(n=15)** | **mRNA-Vax^3^  + BA.1**  **(n=14)** | **mRNA-Vax^3^**  **+ BA.2**  **(n=19)** | **mRNA-Vax^3^  + BA.4/BA.5**  **(n=17)** |
| --- | --- | --- | --- | --- | --- |
| Sex, n (%) |  |  |  |  |  |
| Male | 9 (50) | 9 (60) | 11 (79) | 7 (37) | 7 (41) |
| Female | 9 (50) | 6 (40) | 3 (21) | 12 (63) | 10 (59) |
| Age, median (range) | 38 (23-54) | 69 (61-75) | 32 (23-60) | 30 (25-74) | 33 (22-66) |
| Age group at vaccination, n (%) |  |  |  |  |  |
| 18-55 yrs | 18 (100) | 0 (0) | 12 (86) | 15 (79) | 15 (88) |
| 56-85 yrs | 0 (0) | 15 (100) | 2 (14) | 4 (21) | 2 (12) |
| SARS-CoV-2 status, n (%) |  |  |  |  |  |
| Positive | 0 (0) | 0 (0) | 14 (100)* | 19 (100)** | 17 (100)*** |
| Negative | 18 (100)† | 15 (100)† | 0 (0) | 0 (0) | 0 (0) |
| Unknown | 0 (0) | 0 (0) | 0 (0) | 0 (0) | 0 (0) |
| Interval, median (range) |  |  |  |  |  |
| Days between D1/D2 | ‡ | ‡ | 38 (20-92) | 42 (15-43) | 35 (21-45) |
| Days between D2/D3 | 202 (181-266) | ‡ | 192 (154-256) | 184 (152-259) | 193 (156-305) |
| Days between D3/D4 | N/A | # | N/A | N/A | N/A |
| Days until serum draw after last dose | 28 (26-30) | § | N/A | N/A | N/A |
| Days between last dose/infection | N/A | N/A | 25 (3-112) | 141 (36-200) | 201 (28-263) |
| Days until serum draw after infection | N/A | N/A | 43 (25-55) | 43 (28-99) | 36 (20-58) |

N/A, not applicable; D, dose; yrs, years; n, number.

*, Omicron infection PCR-confirmed at time of recruitment to the research study. Individuals experienced SARS-CoV-2 breakthrough infections between November 2021 and January 2022, during which period the BA.1 lineage was dominant in Germany

**, Individuals experienced SARS-CoV-2 breakthrough infections between March and May 2022, during which period the BA.2 lineage was dominant in Germany. In two cases, BA.2 variant infection was confirmed by sequencing.

***, 5 cases were sequence verified BA.5 infections, 2 cases were family members of individuals with sequence-verified BA.5 infections, 10 individuals experienced SARS-CoV-2 breakthrough infection between June-July 2022, at which time the BA.4 or BA.5 lineage was dominant in Germany

†, No previous clinical or microbiological diagnosis of COVID-19 or positive antigen test for SARS-CoV-2 at baseline

‡, Participants received the BNT162b2 vaccine as part of a governmental vaccination program and the interval between doses was not recorded

#, Participants have received the fourth dose ≥4 months after the third vaccination

§, Serum draw was performed between 28 to 35 Days after vaccination as per protocol

**Table S5. pVN_50_ values of sera collected from SARS-CoV-2-naïve triple-vaccinated individuals (BNT162b2^3^)**

| Participant ID | pVN_50_ | | | | | | |
| --- | --- | --- | --- | --- | --- | --- | --- |
|  | Wild-type | Omi BA.4/5 | Omi BA.4.6/BF.7 | Omi BQ.1.1 | Omi BA.2.75 | Omi BA.2.75.2 | Omi XBB |
| 1 | 160 | 40 | 20 | 10 | 80 | 10 | 5 |
| 2 | 640 | 40 | 40 | 10 | 160 | 20 | 10 |
| 3 | 5120 | 640 | 640 | 320 | 640 | 5 | 40 |
| 4 | 320 | 40 | 20 | 5 | 40 | 20 | 5 |
| 5 | 640 | 40 | 40 | 20 | 40 | 20 | 5 |
| 6 | 320 | 40 | 40 | 20 | 40 | 20 | 5 |
| 7 | 320 | 80 | 40 | 10 | 40 | 20 | 5 |
| 8 | 320 | 80 | 80 | 20 | 80 | 5 | 10 |
| 9 | 160 | 40 | 20 | 5 | 20 | 20 | 5 |
| 10 | 320 | 80 | 40 | 20 | 80 | 5 | 5 |
| 11 | 1280 | 320 | 160 | 40 | 640 | 20 | 40 |
| 12 | 40 | 5 | 5 | 5 | 5 | 20 | 5 |
| 13 | 320 | 40 | 20 | 20 | 80 | 20 | 10 |
| 14 | 160 | 40 | 40 | 10 | 80 | 20 | 5 |
| 15 | 320 | 160 | 80 | 40 | 80 | 160 | 10 |
| 16 | 640 | 80 | 80 | 20 | 80 | 160 | 10 |
| 17 | 2560 | 320 | 160 | 40 | 640 | 40 | 20 |
| 18 | 320 | 80 | 80 | 10 | 80 | 20 | 10 |

**Table S6. pVN_50_ values of sera collected from SARS-CoV-2-naïve quadruple-vaccinated individuals (BNT162b2^4^)**

| Participant ID | pVN_50_ | | | | | | |
| --- | --- | --- | --- | --- | --- | --- | --- |
|  | Wild-type | Omi BA.4/5 | Omi BA.4.6/BF.7 | Omi BQ.1.1 | Omi BA.2.75 | Omi BA.2.75.2 | Omi XBB |
| 19 | 640 | 320 | 160 | 80 | 320 | 40 | 10 |
| 20 | 640 | 160 | 80 | 20 | 80 | 40 | 10 |
| 21 | 1280 | 320 | 80 | 40 | 640 | 80 | 20 |
| 22 | 320 | 160 | 80 | 80 | 320 | 40 | 5 |
| 23 | 640 | 80 | 5 | 5 | 20 | 5 | 5 |
| 24 | 640 | 160 | 40 | 10 | 80 | 20 | 10 |
| 25 | 320 | 40 | 20 | 10 | 160 | 10 | 5 |
| 26 | 1280 | 320 | 160 | 20 | 160 | 80 | 10 |
| 27 | 1280 | 320 | 160 | 20 | 640 | 40 | 20 |
| 28 | 640 | 20 | 5 | 5 | 20 | 5 | 5 |
| 29 | 1280 | 640 | 640 | 80 | 640 | 80 | 40 |
| 30 | 320 | 80 | 40 | 10 | 40 | 10 | 5 |
| 31 | 320 | 40 | 40 | 20 | 160 | 20 | 10 |
| 32 | 1280 | 40 | 40 | 40 | 320 | 40 | 20 |
| 33 | 320 | 80 | 80 | 20 | 40 | 10 | 10 |

**Table S7. pVN_50_ values of sera collected from individuals with Omicron BA.1 breakthrough infection (mRNA-Vax^3^ + BA.1)**

| Participant ID | pVN_50_ | | | | | | |
| --- | --- | --- | --- | --- | --- | --- | --- |
|  | Wild-type | Omi BA.4/5 | Omi BA.4.6/BF.7 | Omi BQ.1.1 | Omi BA.2.75 | Omi BA.2.75.2 | Omi XBB |
| 34 | 1920 | 320 | 640 | 40 | 1280 | 160 | 20 |
| 35 | 960 | 160 | 160 | 20 | 640 | 80 | 10 |
| 36 | 3840 | 640 | 640 | 80 | 640 | 160 | 40 |
| 37 | 960 | 160 | 160 | 40 | 640 | 40 | 40 |
| 38 | 480 | 40 | 80 | 10 | 320 | 20 | 10 |
| 39 | 1920 | 320 | 320 | 40 | 640 | 160 | 40 |
| 40 | 960 | 80 | 160 | 20 | 320 | 40 | 20 |
| 41 | 960 | 80 | 160 | 10 | 160 | 40 | 10 |
| 42 | 480 | 80 | 80 | 20 | 320 | 40 | 20 |
| 43 | 1920 | 2560 | 1280 | 80 | 1280 | 320 | 80 |
| 44 | 3840 | 2560 | 1280 | N/A | 2560 | 640 | N/A |
| 45 | 7680 | 1280 | 1280 | 320 | 5120 | 640 | 160 |
| 46 | 480 | 40 | 40 | 40 | 20 | 10 | 5 |
| 47 | 1920 | 640 | 320 | 80 | 320 | 80 | 40 |

**Table S8. pVN_50_ values of sera collected from individuals with Omicron BA.2 breakthrough infection (mRNA-Vax^3^ + BA.2)**

| Participant ID | pVN_50_ | | | | | | |
| --- | --- | --- | --- | --- | --- | --- | --- |
|  | Wild-type | Omi BA.4/5 | Omi BA.4.6/BF.7 | Omi BQ.1.1 | Omi BA.2.75 | Omi BA.2.75.2 | Omi XBB |
| 48 | 480 | 120 | 40 | 20 | 160 | 20 | 5 |
| 49 | 3840 | 960 | 1280 | 1280 | 2560 | 40 | 40 |
| 50 | 480 | 120 | 80 | 20 | 80 | 20 | 10 |
| 51 | 240 | 120 | 40 | 20 | 40 | 10 | 5 |
| 52 | 1920 | 240 | 160 | 80 | 320 | 40 | 20 |
| 53 | 480 | 240 | 80 | 20 | 160 | 20 | 10 |
| 54 | 960 | 480 | 320 | 160 | 320 | 80 | 20 |
| 55 | 960 | 480 | 320 | 160 | 320 | 160 | 40 |
| 56 | 1920 | 480 | 640 | 320 | 640 | 160 | 80 |
| 57 | 960 | 120 | 80 | 40 | 320 | 40 | 20 |
| 58 | 960 | 480 | 160 | 80 | 320 | 80 | 10 |
| 59 | 1920 | 1920 | 1280 | 640 | 320 | 40 | 40 |
| 60 | 960 | 1920 | 640 | 160 | 320 | 80 | 40 |
| 61 | 240 | 120 | 80 | 40 | 80 | 10 | 5 |
| 62 | 960 | 960 | 640 | 160 | 320 | 80 | 20 |
| 63 | 3840 | 480 | 320 | 160 | 1280 | 80 | 20 |
| 64 | 960 | 240 | 160 | 80 | 160 | 40 | 10 |
| 65 | 960 | 960 | 640 | 160 | 640 | 80 | 20 |
| 66 | 3840 | 480 | 640 | 160 | 1280 | 160 | 40 |

**Table S9. pVN_50_ values of sera collected from individuals with Omicron BA.4/BA.5 breakthrough infection (mRNA-Vax^3^ + BA.4/BA.5)**

| Participant ID | pVN_50_ | | | | | | |
| --- | --- | --- | --- | --- | --- | --- | --- |
|  | Wild-type | Omi BA.4/5 | Omi BA.4.6/BF.7 | Omi BQ.1.1 | Omi BA.2.75 | Omi BA.2.75.2 | Omi XBB |
| 67 | 480 | 480 | 160 | 80 | 160 | 40 | 20 |
| 68 | 1920 | 960 | 1280 | 320 | 160 | 80 | 40 |
| 69 | 960 | 480 | 640 | 160 | 160 | 80 | 40 |
| 70 | 960 | 960 | 640 | 160 | 320 | 40 | 40 |
| 71 | 240 | 60 | 80 | 40 | 80 | 20 | 10 |
| 72 | 1920 | 480 | 640 | 160 | 640 | 80 | 80 |
| 73 | 960 | 240 | 640 | 320 | 160 | 40 | 40 |
| 74 | 960 | 240 | 80 | 40 | 80 | 5 | 5 |
| 75 | 3840 | 1920 | 5120 | 1280 | 2560 | 640 | 320 |
| 76 | 960 | 480 | 640 | 320 | 320 | 80 | 20 |
| 77 | 1920 | 960 | 640 | 320 | 640 | 80 | 40 |
| 78 | 960 | 960 | 320 | 160 | 320 | 80 | 20 |
| 79 | 960 | 960 | 640 | 160 | 640 | 80 | 80 |
| 80 | 960 | 240 | 320 | 320 | 160 | 40 | 20 |
| 81 | 3840 | 3840 | 2560 | 320 | 2560 | 640 | 320 |
| 82 | 480 | 60 | 160 | 40 | 80 | 20 | 10 |
| 83 | 960 | 960 | 80 | 20 | 320 | 20 | 10 |
